# *OsGGCT1* provides tolerance to *Fusarium oxysporum* in *Arabidopsis thaliana* by upregulating γ-glutamyl cycle

**DOI:** 10.64898/2026.05.15.725392

**Authors:** Divya Chaudhary, Richa Vaishnav, Bhoopander Giri, Naveen Chandra Joshi

## Abstract

γ-Glutamyl cyclotransferases (GGCTs) belongs to class of cytosolic enzymes that are responsible for glutathione (GSH) degradation under stress conditions. They regulate GSH homeostasis through the γ-glutamyl cycle which is responsible for maintaining the synthesis of GSH as well as its breakdown, enabling recycling of its constituent amino acids. Although GGCTs have been implicated in enhancing heavy metal (HMs) tolerance in plants, their role in biotic stress remains largely unexplored. Previously, *OsGGCT1* was identified as a gene strongly upregulated in *Fusarium* stress. In this study, the *GGCT1* homolog from *Oryza sativa japonica* was characterized for its role in conferring tolerance to *Fusarium oxysporum* (F.O.). Similar to abiotic factors, biotic stresses significantly impact crop yield and productivity. The rhizosphere harbors diverse microbial communities, including harmful pathogens such as *F. oxysporum*. *Fusarium* causes wilt disease in a variety of plant species, such as: tomato, legumes, rice, and *Arabidopsis thaliana*. Our results demonstrate that overexpression of *OsGGCT1* enhanced tolerance to *F. oxysporum* in *A. thaliana*, primarily by reducing fungal spore accumulation. Transgenic plants showed elevated expression of *OsGGCT1* along with *AtGSH1* and *AtGSH2*, reduced levels of reactive oxygen species (ROS), improved growth and photosynthetic performance and enhanced activities of the antioxidant enzymes. *OsGGCT1* serves as a key component in maintaining GSH homeostasis by supporting glutamate (Glu) regeneration necessary for sustained GSH biosynthesis. Overall, these findings identify *OsGGCT1* as an important constituent of the GSH-mediated detoxification pathway against *Fusarium oxysporum* and provide valuable molecular insights for developing *Fusarium*-tolerant rice varieties with reduced fungal accumulation.

## 1. Introduction

The rise of aggressive, invasive pathogens is a major menace to food security and sustainable agriculture, as they significantly reduce both crop yield and quality. Rice (*Oryza sativa japonica*), a key staple, is among the most widely consumed cereal crops worldwide (Mohidem et al., 2022). Rice yield is affected approximately by 50 different biotic agents like fungi, virus, bacteria, insects, and nematodes. *Fusarium* species are majorly hemibiotroph and are the reason for the mycotoxigenic plant diseases globally (Zakaria, 2023). *F. oxysporum* is a common pathogenic fungus that targets the vascular tissue and causes wilt and rotting of root, collectively known as *Fusarium* wilt. *F. oxysporum* is an ascomycete fungus that inhabitats soil and reproduces asexually. It begins its life cycle as biotroph (1- 5 days) and after 6 days onwards, becomes necrotroph for the later stages of infection (Gordon, 2017). It starts its infection in healthy plants through germinating spores or its mycelia that penetrates the lateral roots, root wounds, or root tips of the plants. In root apoplast, the fungus grows and then reaches the vasculature and colonizes the xylem and drains nutrients and water from the plant that ultimately leads to plant death and release of fungal spores into the soil (Wang et al., 2022). Pattern recognition receptors (PRRs) present on plasma membrane of the host plants can recognize pathogen associated molecular patterns (PAMPs) and damage associated molecular patterns (DAMPs) produced by pathogens and activate pathogen-triggered immunity (PTI) (Chaudhary et al., 2025). To combat diverse stresses, plants generate ROS. This apoplastic ROS burst signals immune activation, serving both as signaling molecules and as toxic agents against pathogens (Waszczak et al., 2018). But the elevated ROS are also harmful to host as well and can cause DNA, RNA, and protein damage. Therefore, plants stimulate the antioxidants like GSH in response to elevated ROS levels. GSH (γ-glutamylcysteinyl-glycine) is a tripeptide that holds a great importance in the living cell. It is present in abundance in eukaryotes. GSH has myriads of functions like resistance to pathogen infection, iron metabolism, scavenging of ROS, antioxidant response, heavy metal detoxification, xenobiotic remediation, etc. (Chaudhary et al., 2024). GSH tripeptide consist of glycine (Gly), cysteine (Cys), and glutamate (Glu), has low-molecular-weight, is non-protein in nature and is widely abundant in aerobic organisms (Dominko and Đikić, 2018). Plants maintain the GSH homeostasis by γ-glutamyl cycle which comprises of biosynthesis of GSH as well as its degradation. The GSH synthesis is a two-step process that requires ATP where *GSH1* carryout the conversion of the Glu, Cys into γ-glutamyl-cysteine and then Gly is added to it by *GSH2* leading to the GSH biosynthesis. The GSH is stored in its oxidised form GSSG in the apoplast. The GSH is degraded into Cys-Gly and 5-OP by the action of GGCTs. 5 oxoprolinase (5- OPase) converts 5-OP into Glu by ATP-dependent reaction. Dipeptidase cleaves the Cys-Gly into constituent amino acid. In *A. thaliana*, GGCTs have three homologs: *GGCT2;1, GGCT2;2* and *GGCT2;3* (Chaudhary et al., 2026). In *Camelina sativa*, the overexpression of the *CsGGCT2;1* gene enhanced resistance to arsenite (As III), with transgenic plants exhibiting greater biomass, higher chlorophyll (Chl) content, and reduced arsenic accumulation compared to wild type (WT) (Singh et al., 2024). In the same way, *GGCT2;1* overexpression enhanced tolerance to As III in *A. thaliana*, resulting in reduced arsenic accumulation and increased biomass compared to the WT. (Paulose et al, 2013). However, role of GGCTs in biotic stresses have not been studied yet. Specifically, our research intended (i) to study the functional role of *OsGGCT1* in conferring *F. oxysporum* stress tolerance by heterologous overexpression in *A. thaliana* and evaluating its impact on plant growth and physiological performance, (ii) to elucidate the involvement of GSH metabolism in stress mitigation, by assessing changes in GSH levels, antioxidant enzyme activities, and ROS accumulation in *OsGGCT1*-overexpressing (OE) line under *F. oxysporum* stress and (iii) to examine the molecular regulation of the GSH biosynthetic pathway under pathogen stress, by analysing the gene expression levels of *AtGSH1* and *AtGSH2* in response to *OsGGCT1* overexpression. Collectively, this study highlights *OsGGCT1* as a significant regulator of GSH-mediated defense, offering a promising genetic target for improving plant resilience against *F. oxysporum* stress.

## 2. Materials and methods

### 2.1. Plant materials, transformation, and screening

From the Rice Genome Annotation Project (RAP) database, *OsGGCT1* gene sequence was obtained and on the basis of the *OsGGCT1* cDNA sequence the gene specific primers were designed (Forward and reverse) (Table 1), and were used to amplify its coding region from rice cDNA, which was then cloned into plant transformation vector pGWB441 which was further cloned in *Agrobacterium tumefaciens* GV3101. This construct was next transformed into *A. thaliana* (Col-0) by vacuum infiltration method. To obtain the homozygous T_3_ generation, the primary transformants were subjected to MS medium- half strength containing 50 mg/L kanamycin and selected. The transgenic lines were confirmed by genomic DNA (gDNA) PCR using 35S internal forward primer (FP), *OsGGCT1* reverse primer (RP), gene specific FP and RP and kanamycin-specific FP and RP.

**Table 1:**
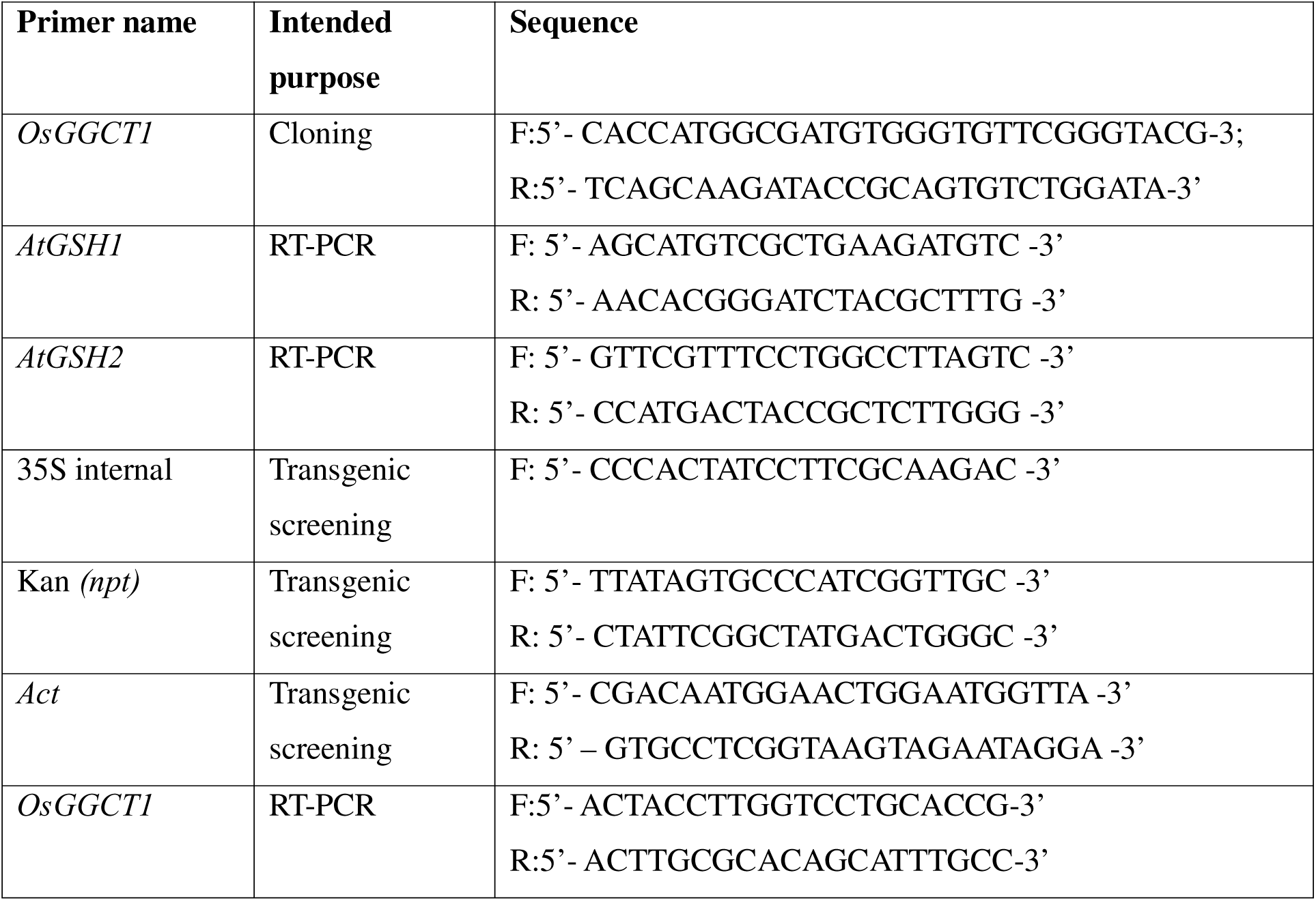
Primers used in the study.

### 2.2. RNA isolation and qRT- PCR to analyse changes in gene expression

Total RNA was isolated from 21-day-old *A. thaliana* seedlings with the use of RNeasy® Plant Mini Kit (Qiagen, Germany). Using the Verso cDNA Synthesis Kit (Thermo Scientific™), first- strand cDNA was synthesized from 2 µg RNA using oligo (dT) primers. 1 µl of cDNA (1: 20 dilution) was used as template and using gene specific primers, qRT- PCR was performed on a Bio-Rad real-time PCR system (Table 1). Analysis of gene expression was done with comparative CT (2^−ΔΔCT) method and normalized against Ubiquitin (*Ubq*) (Schmittgen and Livak, 2008). The data is expressed as mean ± standard error for three biological replicates.

### 2.3. *Fusarium* infection in WT and *OsGGCT1* transgenics

*A. thaliana* (WT and transgenic lines) seeds were sterilized with 90 % ethanol and 70 % ethanol containing Tween 20 and then by repetitive washing by autoclaved distilled water. The *A. thaliana* seeds were germinated on MS medium with 1 % sucrose at 22 °C with a 16 h light/8 h dark photoperiod. Upon reaching the 2-week stage these seedlings were infected with *F. oxysporum* (ITCC 8094) grown in Potato Dextrose Broth (PDB) at 28 °C for 4 days (10c spores). Roots were dipped in the spore suspension and transferred to sucrose-free MS medium, while controls were mock-treated with PDB. After six days, plants showing symptoms were harvested in three biological replicates for further analysis.

### 2.4. Analysis of morphological and physiological parameters

21-day-old WT and *OsGGCT1* transgenic seedlings were evaluated for morphological parameters like fresh weight (FW), dry weight (DW), and root length (RL). Also, the physiological parameters like total Chl, and carotenoid contents were measured. FW, DW and RL were measured directly by ruler, while pigments were extracted from 100 mg tissue using 90 % cold acetone. Centrifugation was performed at 10,000 rpm for 10 min at the temperature of 4 °C. At 663, 645, and 450 nm, the absorbance of the supernatant was noted. According to and Jensen, (1978) and Porra et al. (1989), the carotenoids and total Chl total were calculated respectively.

### 2.5. Estimation of hydrogen peroxide (H_2_O_2_) content

From 21-day-old seedlings of WT and *OsGGCT1* transgenics, a sample of 0.1 g was homogenized in 1 % TCA followed by centrifugation at 12,000 rpm for around 10 min. Then to potassium phosphate buffer (pH 7) and potassium iodide (KI), supernatant was added. The absorbance was recorded at 390 nm. The HcOc levels were calculated using a standard curve (Loreto and Velikova, 2001).

### 2.6. Quantification of GSH content

From 21-day-old seedlings of WT and *OsGGCT1* transgenics, a sample of 0.1 g was crushed in pre-chilled 5 % TCA (1 ml), followed by centrifugation at 10,000 rpm for 10 min. To potassium phosphate buffer (pH 7) having 5,5′-dithiobis-(2-nitrobenzoic acid) (DTNB), supernatant was added and at 412 nm, absorbance was recorded. The levels of GSH were calculated using a standard curve (Tyagi et al., 2018).

### 2.7. Antioxidant enzyme assay

#### 2.7.1. Peroxidase (POD)

100 mg of seedlings were crushed in phosphate buffer (pH 7) containing 1 mM ethylenediaminetetraacetic acid (EDTA) and polyvinyl pyrrolidone (PVP) (2 %), followed by centrifugation at 12,000 rpm for 20 min. 100 µl of enzyme extract was added to a mixture of guaiacol, HcOc, and phosphate buffer (pH 7), and absorbance was measured at 470 nm (Varga et al., 2013).

#### 2.7.2. Catalase (CAT)

100 mg of seedlings were homogenized in phosphate buffer (pH 6.8) followed by centrifugation at 12,000 rpm for 20 min. Reaction was initiated by adding enzyme extract to HcOc-containing buffer (with a control lacking HcOc), and the decline in absorbance at 240 nm was monitored to assess HcOc breakdown (Reddy et al., 2004).

#### 2.7.3. Superoxide dismutase (SOD)

50 μl of crude enzyme extract was added to a reaction mixture containing phosphate buffer (0.1 M) (pH 7.9), EDTA (3 mM), nitro blue tetrazolium (NBT) (2 mM), methionine (200 mM), and riboflavin (75 μM). The mixture was incubated under white light at 25 °C for 15 minutes, and absorbance was recorded at 560 nm (Dhindsa et al., 1981).

### 2.8. In situ detection of H_2_O_2_ and super oxide radical (O_2_ ^-^)

21-day-old leaves of WT and *OsGGCT1* transgenic were used to detect HcOc and Occ through 3,3-diaminobenzidine (DAB) and NBT staining, respectively. Both stressed and control leaves were incubated overnight at room temperature in 0.1 % DAB or 0.2 % NBT solutions, followed by decolorization in absolute ethanol by boiling for 15 minutes (Kumar et al., 2014).

### 2.9. Percent root colonization

Percent root colonization was performed using Phillips and Hayman, (1970) method. Roots of 21-day-old WT and *OsGGCT1* transgenic were assessed for colonization. Roots were first washed with distilled water, followed by 10 % potassium hydroxide (KOH) treatment for 15 min then with 1 M hydrochloric acid (HCl) for 10 min, and finally stained overnight with 0.02% trypan blue. Before observation, roots were destained in 50% lactophenol. Random 1 cm root segments (20 per treatment) were examined under a light microscope. Colonization (%) was calculated as: (colonized segments / total segments examined) × 100.

### 2.10. Statistical analysis

The experiments were performed in triplicates, and results are expressed as mean ± standard error. Statistical significance was evaluated using one-way and two-way ANOVA, with *p* ≤ 0.05 considered significant. Significance levels are denoted as *, **, ***, and **** for *p* ≤ 0.05, 0.01, 0.001, and 0.0001, respectively. Data were analyzed using GraphPad Prism (v8.0.2).

## 3. Results

### 3.1. Development and characterization of transgenic lines in *Arabidopsis*

A construct harbouring *OsGGCT1* cDNA in the sense (overexpression) orientation under the constitutive expression of CaMV 35S promoter was introduced into *A. thaliana* (Col-0) via *Agrobacterium* mediated transformation (**Figure 1 A**) and more than 17 transgenic lines were generated. From each transformed plant, the seeds were collected and stored. On half-strength MS medium supplemented with 50 mg/L kanamycin, the seeds were screened. All lines exhibited a 3:1 segregation ratio on the selection plates, and the surviving seedlings were advanced to the T_2_ generation and further selected for kanamycin resistance. Homozygous T_3_ transgenic lines were subsequently confirmed by gDNA PCR (**Figure 1 B-D**). Gene-specific FP and RP, a 35S promoter–specific FP paired with a gene-specific RP, and kanamycin-specific FP and RP were used.

**Figure 1:**
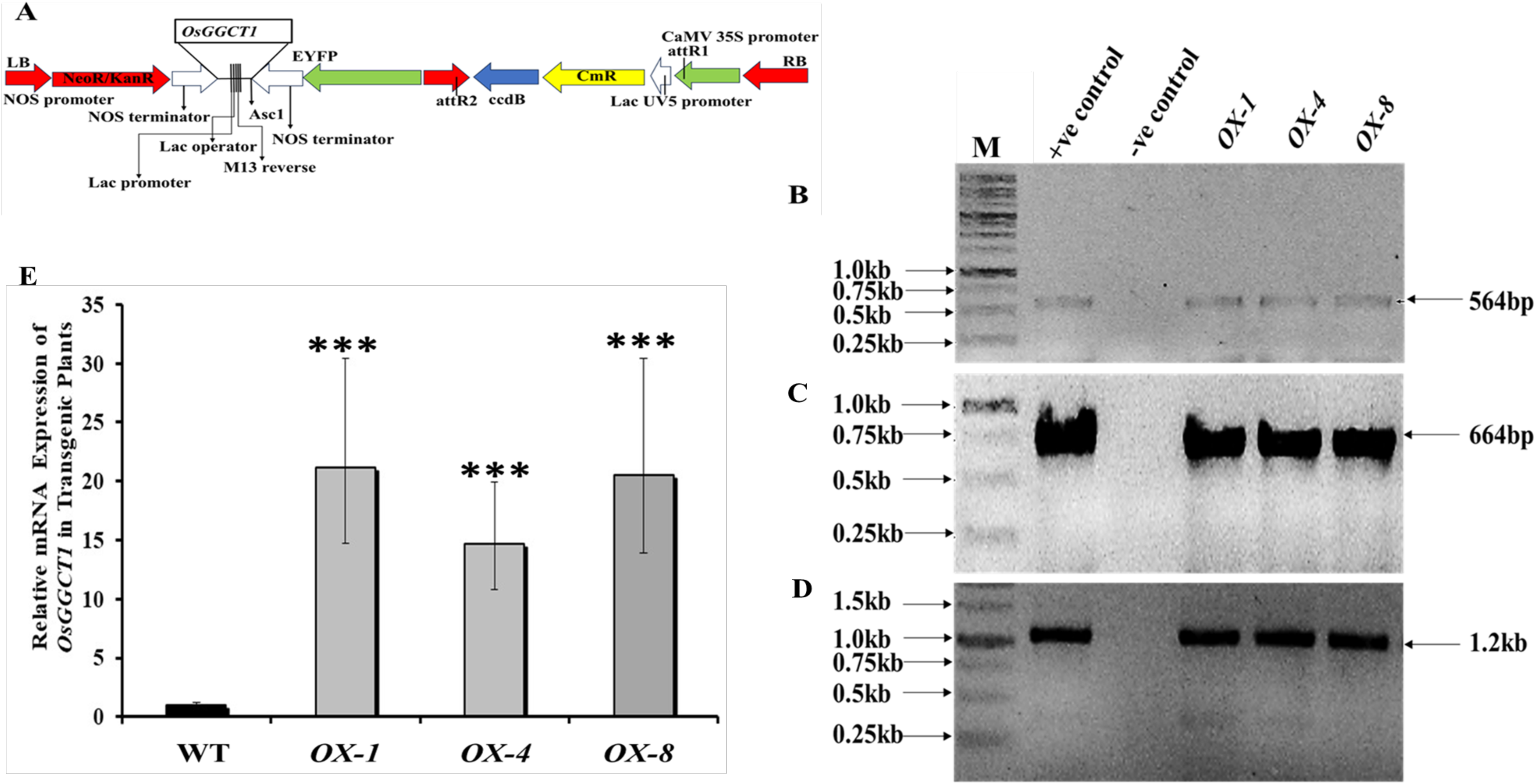
Transformation of *Arabidopsis* by *OsGGCT1* cDNA. *OsGGCT1* cloned in pGWB441 modified plant transformation vector was used to transform *Arabidopsis* plants. (A) vector construct, (B-D) confirmation of T_3_ generation of *OsGGCT1* transgenic line by PCR using gDNA as template and gene specific forward and reverse primers, 35Sforward primer and gene specific reverse primer, and kanamycin specific forward and reverse primers, (E) expression of *OsGGCT1* cDNA in transgenic over-expresser (*OX- 1,4* and *8*) by quantitative real time PCR. Standard deviation is shown by error bar, \**p* ≤ 0.05, \*\**p* ≤ 0.01, and \*\*\**p* ≤ 0.001 are the stated significance levels.

Following confirmation of the transgenic lines by gDNA PCR, the expression of *OsGGCT1* in both transgenic plants and WT was assessed using qRT-PCR. For this analysis, 1 µg of RNA extracted from WT and *OsGGCT1* transgenic plants was used to synthesize cDNA, which was then subjected to qRT-PCR. Out of 17, 3 lines were selected which showed the highest expression of *OsGGCT1* by qRT-PCR. The analysis revealed that the *OsGGCT1* OE lines (*OX-1,4,* and *8*) had 15–20-fold increase in the gene expression (**Figure 1 E**). On the basis of the qRT-PCR gene expression analysis the highest expression was showed by *OX-8*, so all the experiments under *F. oxysporum* stress were carried out using WT and *OX-8* line.

### 3.2. Analysis of morphological and physiological parameters

After 21 days of growing the WT and *OsGGCT1* OE lines (*OX-1, OX-4*, and *OX-8*) under normal (untreated) conditions, the morphological and physiological parameters were analysed. Under normal conditions, the phenotype of OE lines in comparison to WT was non-significant (**Supplementary figure 1**). The difference in RL, FW, and DW observed between OE lines and WT was non- significant (**Supplementary figure 2 A, B, C**). Also, the difference in physiological parameters like total Chl and carotenoids observed between WT and OE lines were non- significant (**Supplementary figure 2D, E**).

### 3.3. Analysis of GSH and H_2_O_2_ content

After 21 days of growing the WT and *OsGGCT1* OE lines (*OX-1, OX-4*, and *OX-8*) under normal (untreated) conditions, the activities of GSH and H_2_O_2_ were assessed. No significant changes in GSH and H_2_O_2_ levels were observed between OE lines and WT under normal conditions **(Supplementary figure 3 A, B).**

### 3.4. Analysis of enzyme activities

After 21 days of growing the WT and *OsGGCT1* OE lines (*OX-1, OX-4*, and *OX-8*) under normal (untreated) conditions, the enzyme activities like CAT, SOD and POD were assessed. Under normal conditions, the differences observed between WT and OE lines were non-significant **(Supplementary figure 4 A, B, C).**

### 3.5. Analysis of morphological and physiological parameters of the WT and *OX-8* under *F. oxysporum* stress

After 21 days of growing the WT and *OX-8* under normal (untreated) and *F. oxysporum* stress, the morphological and physiological parameters were analysed. Under untreated conditions, the differences observed in any of the parameter between WT and *OX-8* were non- significant. However, in the presence of *F. oxysporum* treatment, *OX-8* exhibited strong tolerance and produced greater shoot and root biomass compared to WT plants **(Figure 2).** The *OX-8* had higher FW, DW, and RL than the WT under *F. oxysporum* stress **(Figure 3 A, B, C)**. The FW, DW and RL decreased by 78.9 %, 63.86 % and 69.10 % respectively in WT under *F. oxysporum* stress, whereas, in *OX-8*, the FW, DW and RL decreased only by 33.8 %, 46.30 % and 42.10 % respectively. The total Chl and carotenoids contents of *OX-8* were also significantly higher than the WT under *F. oxysporum* stress **(Figure 3 D, E).** The total Chl and carotenoid content decreased by 38.92 % and 54.87 % respectively in WT under *F. oxysporum* stress, whereas, in *OX-8*, the total Chl and carotenoid content decreased only by 17.21 % and 29.85 % respectively.

**Figure 2:**
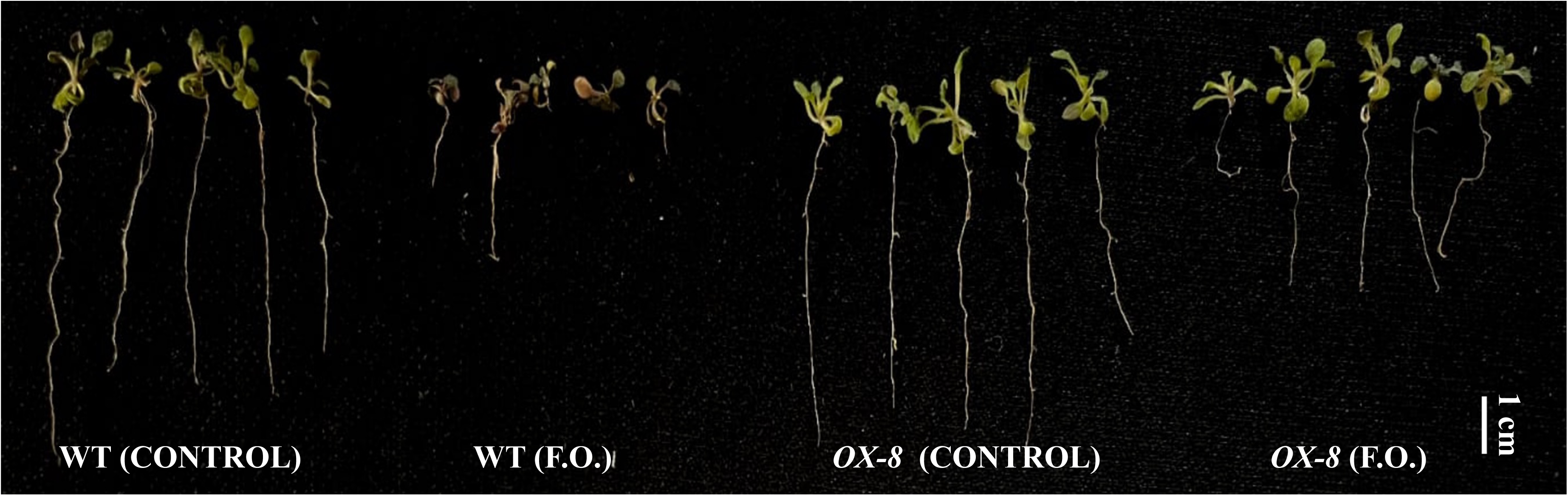
Phenotype of WT and *OX-8* under *Fusarium* stress. WT and transgenic line (*OX-8*) were grown on MS medium under normal (without *Fusarium*) and *Fusarium oxysporum* (F.O.) conditions at 22 c under 16h/8h light/dark cycle for 21 days and their phenotype was analyzed.

**Figure 3:**
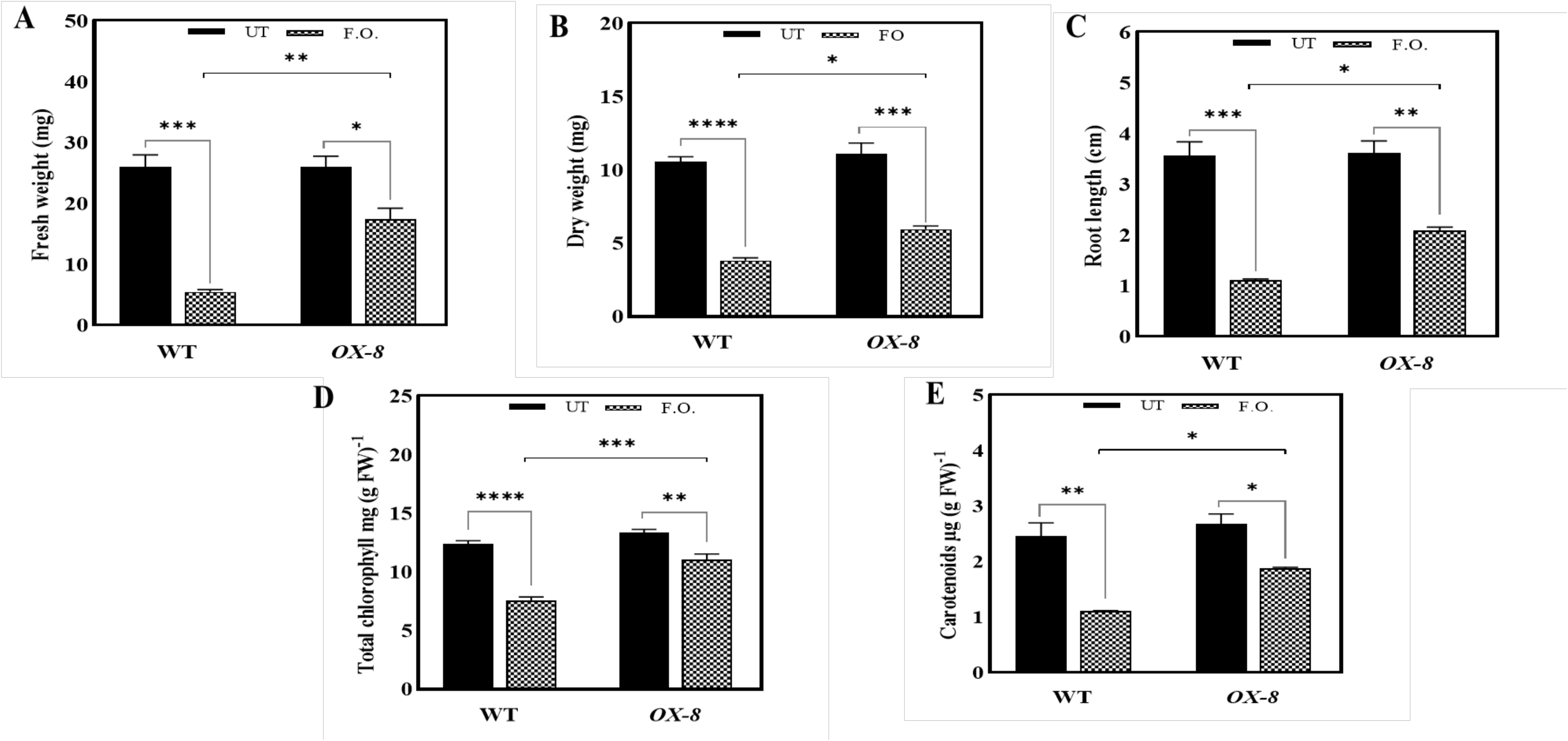
Morphological and physiological parameters of WT and *OX-8* under *Fusarium* stress. WT and transgenic line (*OX-8*) were grown on MS medium under normal (without *Fusarium*) and *Fusarium oxysporum* (F.O.) conditions at 22 c under 16h/8h light/dark cycle for 21 days and their morphological and physiological parameters were analyzed. (A) fresh weight, (B) dry weight, (C) root length, (D) total Chl and (E) carotenoid content. The tests were analyzed using two-way ANOVA, where *, **, and *** represent significance and **** high significance at *p*c≤c0.05, 0.01, 0.001, and 0.0001, respectively.

### 3.6. Analysis of GSH and H_2_O_2_ content of the WT and *OX-8* under *F. oxysporum* stress

After 21 days, GSH and H_2_O_2_ levels were evaluated in WT and *OX-8* under control and *F. oxysporum* stress conditions, with no differences observed between them under normal conditions. In contrast, *F. oxysporum* led to a reduction in GSH levels in WT **(Figure 4A)**, while HcOc levels were markedly higher in WT than in *OX-8* **(Figure 4 B)**. The GSH content decreased by 57.89 % in WT whereas, in *OX-8*, the GSH content decreased only by 31.55 %. The H_2_O_2_ content increased by 52.9 % in WT whereas, in *OX-8*, the H_2_O_2_ content increased only by 10.28 % reflecting the better elimination H_2_O_2_ by transgenic line.

**Figure 4:**
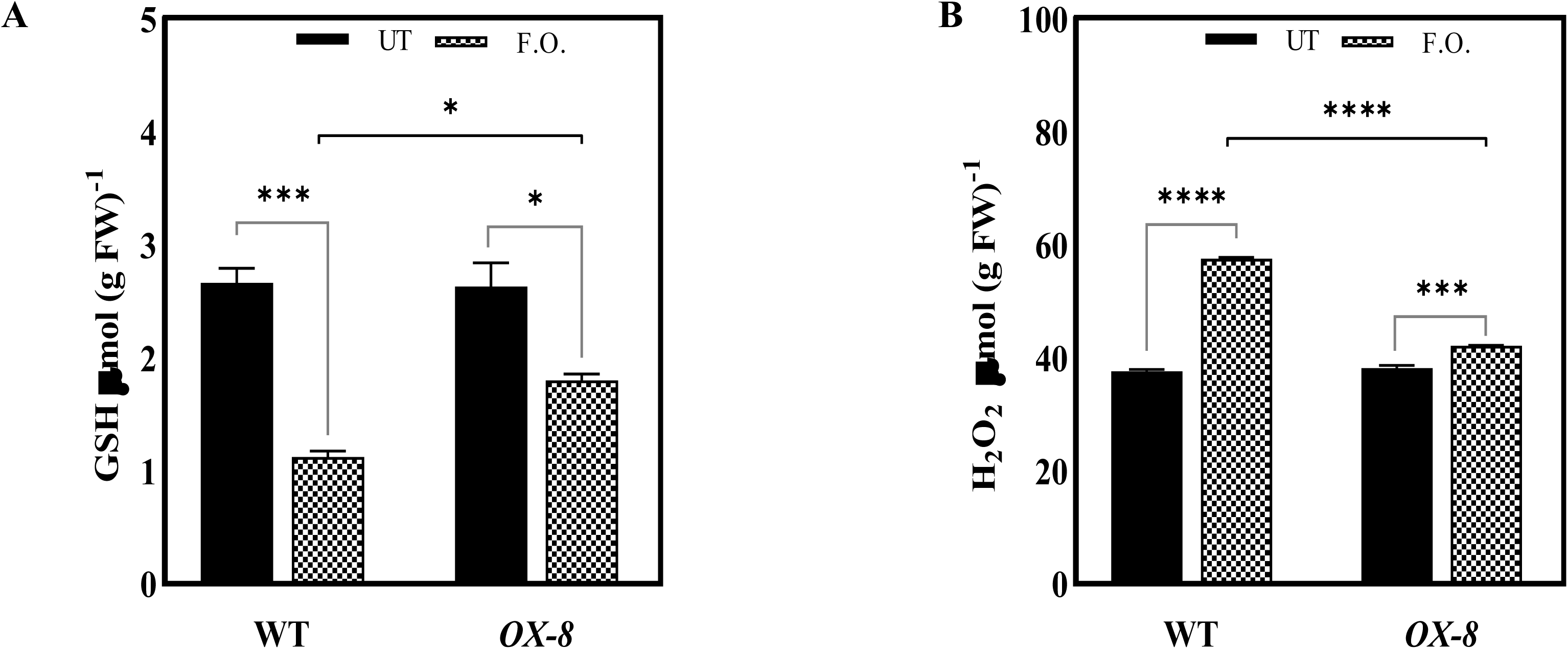
GSH and H_2_O_2_ levels of WT and *OX-8* under *Fusarium* stress. WT and transgenic line (*OX-8*) were grown on MS medium under normal (without *Fusarium*) and *Fusarium oxysporum* (F.O.) conditions at 22 c under 16h/8h light/dark cycle for 21 days and their (A) GSH (B) H_2_O_2_ contents were analyzed. The tests were analyzed using two-way ANOVA, where *, **, and *** represent significance and **** high significance at *p*c≤c0.05, 0.01, 0.001, and 0.0001, respectively.

### 3.7. Analysis of enzyme activities in the WT and *OX-8* under *F. oxysporum* stress

Under both normal conditions and after *F. oxysporum* treatment in MS medium, the enzyme activities were assessed in WT and *OX-8* (**Figure 5 A, B, C).** *F. oxysporum* treatment increased the activities of POD, CAT and SOD enzymes in both WT and *OX-8*; however, the induction was more pronounced in the transgenic line, resulting in higher antioxidant enzyme activities in *OX-8* compared to WT. These findings suggest that the reduced oxidative stress observed in *OX-8* is likely due to its elevated antioxidative enzyme activity under *F. oxysporum* stress relative to the WT. The levels of CAT, POD and SOD increased by 98.08 %, 26.60 % and 129.6 % respectively in WT under *F. oxysporum* stress, whereas, the levels of CAT, POD and SOD increased by 235.9 %, 95.83 % and 386.5 % respectively in *OX-8*.

**Figure 5:**
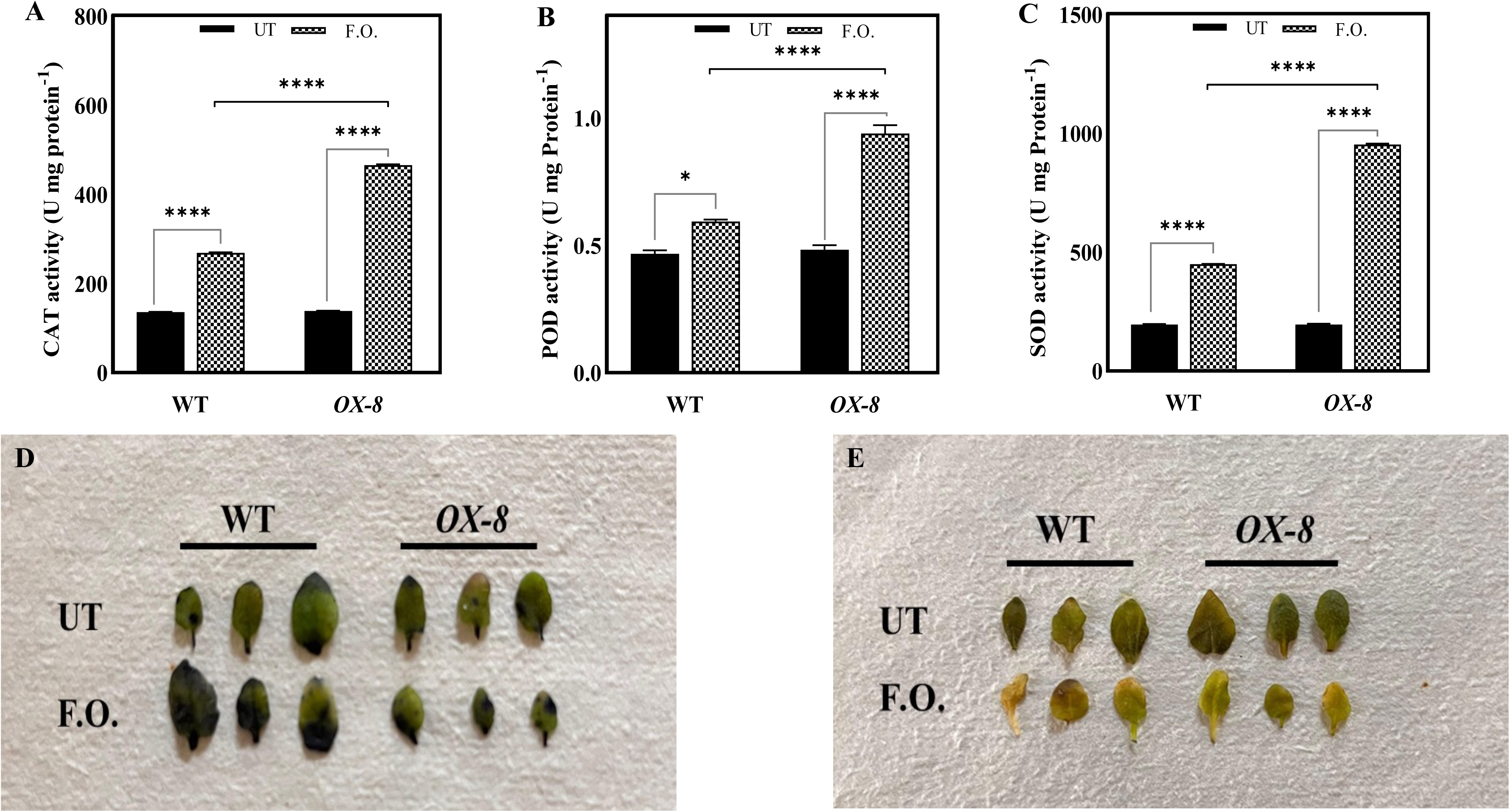
Enzyme activities and NBT and DAB histochemical staining of WT and *OX-8* under *Fusarium* stress. WT and transgenic line (*OX-8*) were grown on MS medium under normal (without *Fusarium*) and *Fusarium oxysporum* (F.O.) conditions at 22 c under 16h/8h light/dark cycle for 21 days and their enzyme activities were analyzed. (A) CAT, (B) POD and (C) SOD. Their leaves were stained with (A) NBT and (B) DAB for detection of O^2-^, H_2_O_2_ radicals respectively. The tests were analyzed using two-way ANOVA, where *, **, and *** represent significance and **** high significance at *p*c≤c0.05, 0.01, 0.001, and 0.0001, respectively.

### 3.8. *OX-8* exhibited lower oxidative stress as indicated by NBT and DAB staining

*F. oxysporum* enhances oxidative stresses. Visualization of O^2-^, H_2_O_2_ by NBT and DAB staining, respectively, confirmed low levels of O^2-^, H_2_O_2_ in *OX-8* when compared with WT **(Figure 5 D, E)**.

### 3.9. Root colonization in the WT and *OX-8* under *F. oxysporum* treatment

To assess the presence of *F. oxysporum* spores in the root cells of WT and the transgenic *OX-8*-line, root colonization analysis was performed. Microscopic observations revealed that WT plants (42 %) accumulated more fungal spores than *OX-8* (35 %), indicating that the improved *F. oxysporum* tolerance in the transgenic line is associated with lower spore accumulation **(Figure 6 A).**

**Figure 6:**
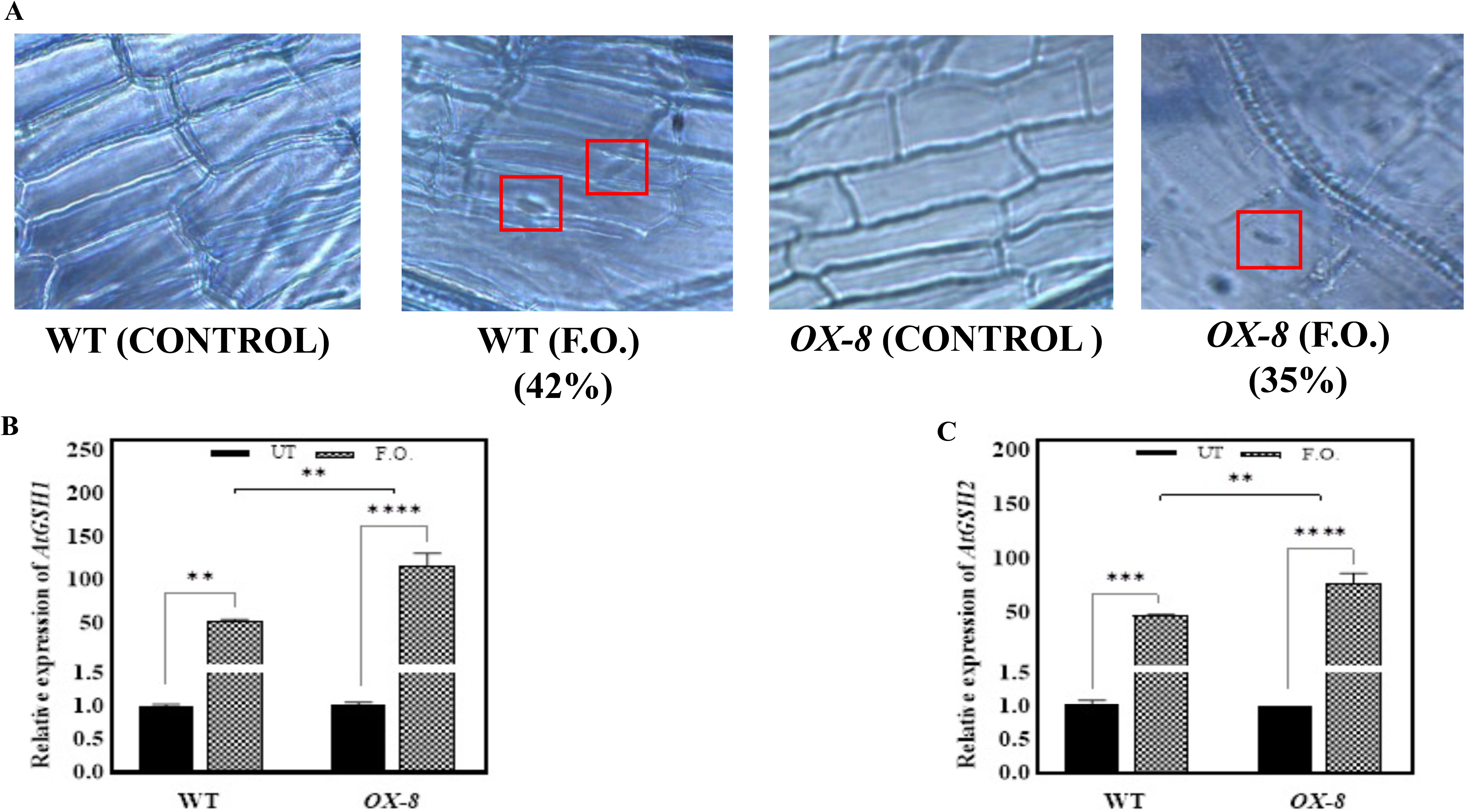
Root colonization and gene expression analysis of GSH biosynthesis genes in WT and *OX-8* under *Fusarium* stress. WT and transgenic line (*OX-8*) were grown on MS medium under normal (without *Fusarium*) and *Fusarium oxysporum* (F.O.) conditions at 22 c under 16h/8h light/dark cycle for 21 days and their root colonization (A) was done. qRT- PCR was performed to analyze the expression of (B) *AtGSH1* and (C) *AtGSH2.* The tests were analyzed using two-way ANOVA, where *, **, and *** represent significance and **** high significance at *p*c≤c0.05, 0.01, 0.001, and 0.0001, respectively.

### 3.10. Expression levels of *AtGSH1* and *AtGSH2* in WT and *OX-8* under *F. oxysporum* stress

To elucidate the mechanism underlying *OX-8* tolerance to *F. oxysporum*, we analyzed the *AtGSH1* and *AtGSH2* expression levels. Both *AtGSH1* and *AtGSH2* genes were upregulated under stress conditions in WT and *OX-8*, with a markedly higher upregulation observed in *OX-8* **(Figure 6 B, C).** The relative expression of *AtGSH1* and *AtGSH2* was upregulated by 444 % and 183 % in WT under *F. oxysporum* stress, whereas, in case of *OX-8*, the relative expression of *AtGSH1* and *AtGSH2* was upregulated by 610 % and 380 % respectively.

## 4. Discussion

*Fusarium* genus contains filamentous fungi that are ascomycetes in nature and are widely distributed in soil (<u>Jackson et al., 2024</u>). *F. oxysporum* is the, most pervasive, anamorphic soil- borne pathogen that significantly contributes to losses of the crops of high economic status. It causes wilt disease on several plant species including rice and *A. thaliana* (Lal et al., 2024). *Fusarium* wilt is characterized by symptoms such as discoloration of the vascular tissues and wilting of older leaves, which gradually progress to tissue necrosis, leaf shedding, and eventually complete plant death (<u>Ahmad et al., 2023</u>). Through the process called conidiation, it produces three types of asexual spores: macroconidia, chlamydospores and microconidia (<u>Ohara and Tsuge, 2004</u>). It’s spores penetrate the host through the roots and the penetration increases by the hydrolytic enzymes secreted by *F. oxysporum*. *Fusarium* mycelium invades the root cortex, breaches the endodermis, and eventually enters the xylem, where it obstructs water transport. In the xylem, it forms microconidia, decreases transpiration, and leads to vascular wilt, root rot, and other disease symptoms in the host plant (Mariyappa et al., 2025). Our results of the root colonization studies showed that the root colonization by *Fusarium* spores were higher than the WT. The % root colonization in WT was 42 % whereas, in *OX-8*, the % root colonization was 35 %. These results showed that the transgenic line *OX-8* showed higher tolerance to *F. oxysporum* stress and plants showed better survival rate and greater biomass as compared to WT.

*F. oxysporum* infection after 7 days in 14-day old WT resulted in loss of the morphological and physiological parameters. The plants appeared to be rotted and the RL, FW and DW decreased by 69.9, 78.9, and 63.86 % respectively. However, the *OX- 8* had higher FW, DW and RL as compared to WT under stress. Under *F. oxysporum* stress, the RL, FW, and DW decreased only by 42.10, 33.28, and 46.30 % respectively. The process by which plants synthesize nutrients is called photosynthesis and has an important role in sustaining life on Earth (<u>Nolasco, 2020</u>). Fungal infection leads to reduced rate of photosynthesis (<u>Yang and Luo, 2021</u>). Reduction of Chl content is characterized by yellowing of leaves. A decrease in photosynthesis and photosynthetic parameters in presence of necrotrophic fungus like *Xylella fastidosa*, *Puccinia psidii*, *Plasmopara viticola* and *Puccinia striformis* has been reported (Cheaib and Killiny, 2025). The total Chl and carotenoid content of WT reduced by 38.92 and 54.87 % respectively under *F. oxysporum* stress condition. In case of *OX-8*, the total Chl and carotenoids decreased only by 17.21 and 29.85 % respectively. This could be due to the induction of defense responses in the transgenic line to fight against pathogens, plants interconnect the photosynthesis to immune defense. Plants may reduce photosynthesis to restrict the supply of nutrients to invading pathogens (Sowden et al., 2018). Fungal infection disturbs plant metabolism, leading to excessive production of ROS like, HcOc, Occ, and nitric oxide (NO•), which ultimately results in oxidative stress (Sahu et al., 2022). The serious damage to biological components is caused by ROS. The levels of H_2_O_2_ increased by 52.9 % in WT whereas in *OX-8* it only increased by 10.28 % under *F. oxysporum* stress. The NBT and DAB staining also showed the same results where the *OX-8* under *F. oxysporum* stress had lower O^2-^ and H_2_O_2_ levels as compared to WT. Sahu et al. (2024), also documented the elevated levels in H_2_O_2_ and enzymatic activities in wheat upon *Fusarium* infection. Plants generate a variety of both enzymatic and non- enzymatic antioxidants in order to control the higher ROS levels (Rudenko et al., 2023). Research suggests that the activities of antioxidant enzymes like POD, CAT, and SOD rise in response to pathogen attack. In our study, we observed that CAT, POD, and SOD levels increased by 98.08 %, 26.60 %, and 129.6 %, respectively, in WT. In *OX-8*, the levels of CAT, POD, and SOD increased by 235.9 %, 95.83 %, and 386.5 %, respectively. The increased activity of these enzymes may result from the induction and upregulation of stress-responsive genes. The GSH is a vital antioxidant with numerous functions; it facilitates the ROS detoxification and helps protect plants from oxidative damage (Iqbal et al., 2025). Therefore, increasing the GSH synthesis can increase the cellular defense against oxidative stress. The pathway of GSH homeostasis is well established, therefore, manipulating the enzymes involved in GSH homeostasis is a good approach to enhance tolerance to oxidative stress (<u>Lushchak, 2012</u>). Decreased GSH levels impairs the defense against pathogens by disrupting its role as redox buffer (Georgiou-Siafis et al., 2023). The levels of GSH decreased by 57.89 % in WT whereas in *OX-8* its levels decreased only by 31.55 % only. We also studied the gene expression levels of GSH biosynthesis genes. The *AtGSH1* and *AtGSH2* gene were upregulated by 444 and 183 % respectively on WT whereas in *OX-8*, the *AtGSH1* and *AtGSH2* gene were upregulated by 610 and 380 % respectively. As a result, GSH biosynthesis is stimulated, along with enhanced recycling of GSH via the γ-glutamyl cycle in the *OX-8* line. (**figure 7).**

**Figure 7:**
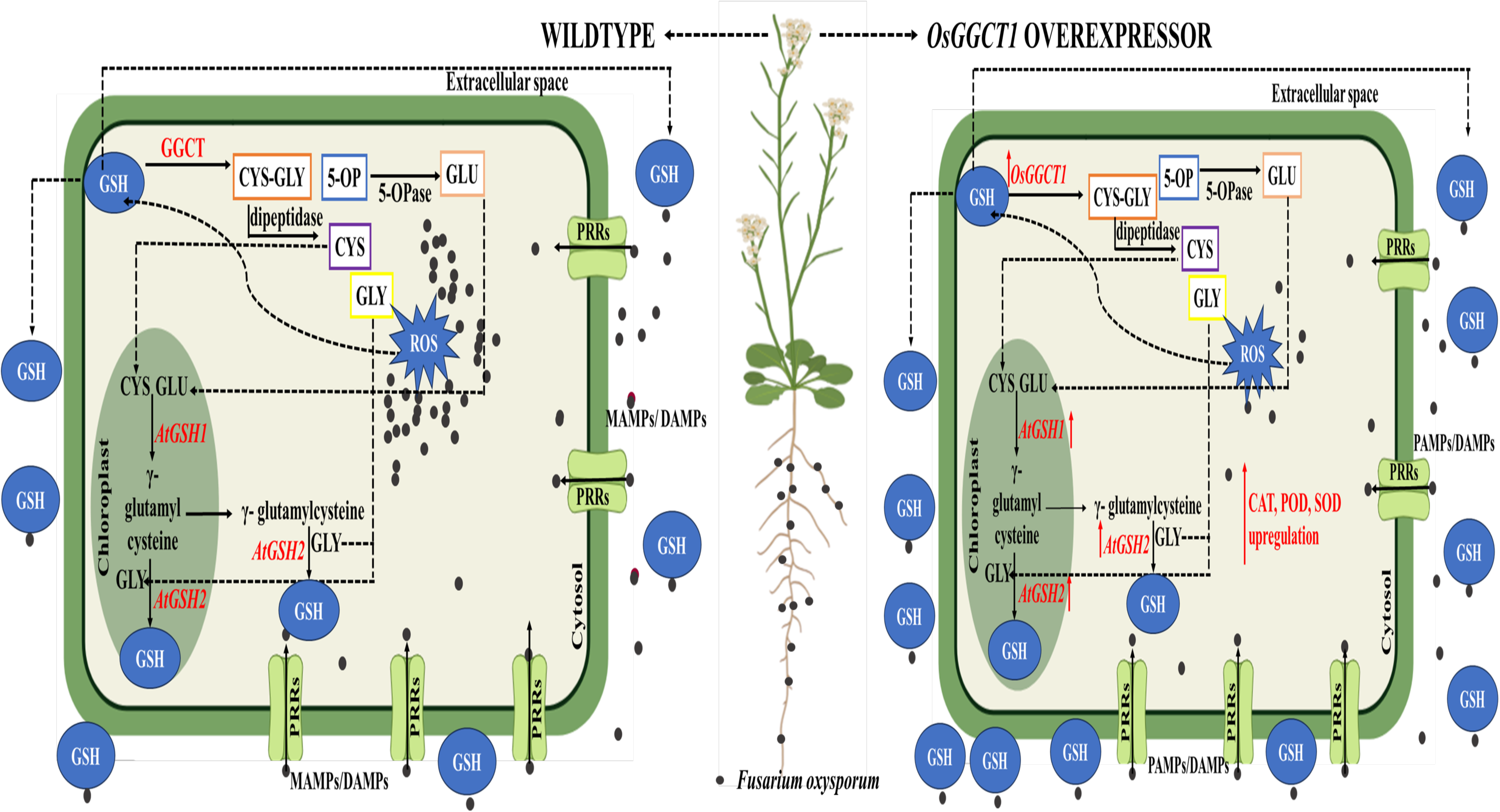
Model of *OsGGCT1* expression in *Arabidopsis thaliana*. Endogenous priming by *OsGGCT1* confers tolerance to *Fusarium* stress. *OsGGCT1* mediates GSH degradation into Cys–Gly and 5- OP (further converted into Glu by 5-OPase), promoting amino acid recycling and stimulating GSH biosynthesis via *AtGSH1* and *AtGSH2*. Enhanced GSH turnover improves redox homeostasis, reduces ROS accumulation, and upregulates antioxidant enzymes (CAT, POD, SOD), thereby conferring increased *Fusarium* tolerance. *Abbreviations*: *OsGGCT1* (rice γ- glutamylcyclotransferase1), GSH (glutathione), Cys (cysteine), Gly (glycine), Glu (glutamate), 5- OP (5-oxoproline), 5-OPase (5-Oxoprolinase), ROS (reactive oxygen species), *AtGSH1* (*Arabidopsis* glutathione synthetase1), *AtGSH2* (*Arabidopsis* glutathione synthetase2), CAT (catalase), POD (peroxidase), and SOD (superoxide dismutase), PRRs (pathogen recognition receptors), PAMPs/ DAMPs (pathogen associated molecular patterns/ damage associated molecular patterns).

## Conclusion

This study shows that overexpressing *OsGGCT1* in *Arabidopsis thaliana* markedly improves the plant’s tolerance to *Fusarium oxysporum* stress. As compared to the *OsGGCT1* OE line (*OX-8*) exhibited improved biomass, reduced fungal colonization, elevated chlorophyll and carotenoid contents, increased glutathione (GSH), antioxidant enzyme activities, and lower reactive oxygen species (ROS) accumulation. Notably, the upregulation of the *AtGSH1* and *AtGSH2* indicates that *OsGGCT1* promotes efficient GSH recycling, thereby strengthening redox homeostasis under stress conditions. Overall, this study highlights the successful functional translation of a rice gene into *Arabidopsis thaliana*, underscoring its potential in improving plant resilience. The findings pave the way for developing *Fusarium*-tolerant crops and phytoremediators capable of sustaining productivity in contaminated or marginal soils, contributing to improved food safety and long-term agricultural sustainability.

## Author contribution

D.C. performed the experiments and compiled the manuscript. R.V. and B.G. helped in the data analysis and N.C.J conceptualized the research.

## Acknowledgement

This research is supported by DST/SERB-SRG file no. SRG/2020/002237.

## Figure legends

**Supplementary figure 1:**
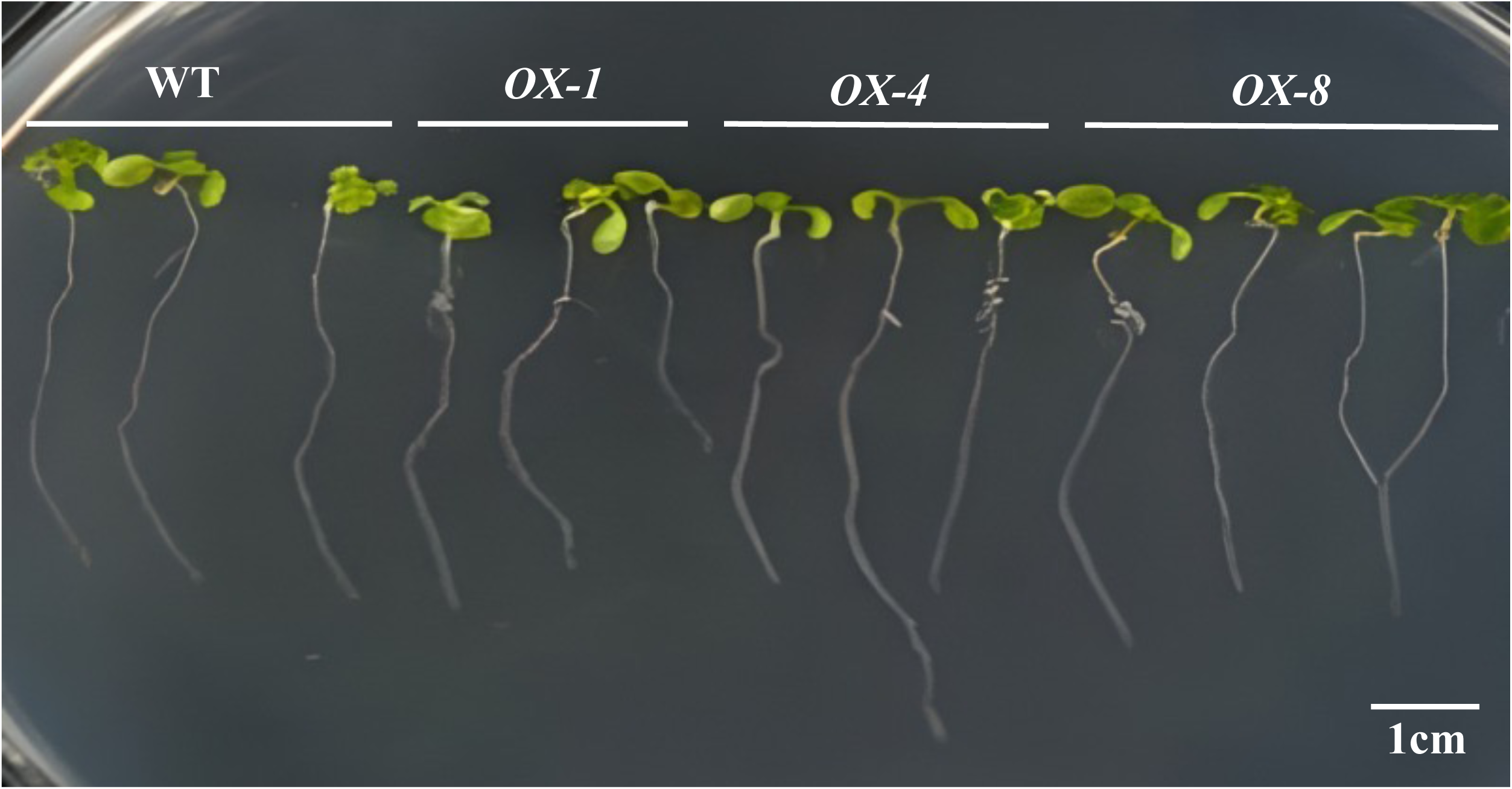
Phenotype of *OsGGCT1* transgenic plants. WT and transgenic lines (*OX- 1,4* and *8*) were grown on MS medium under normal conditions at 22 c under 16h/8h light/dark cycle for 21 days and their phenotype was analyzed.

**Supplementary figure 2:**
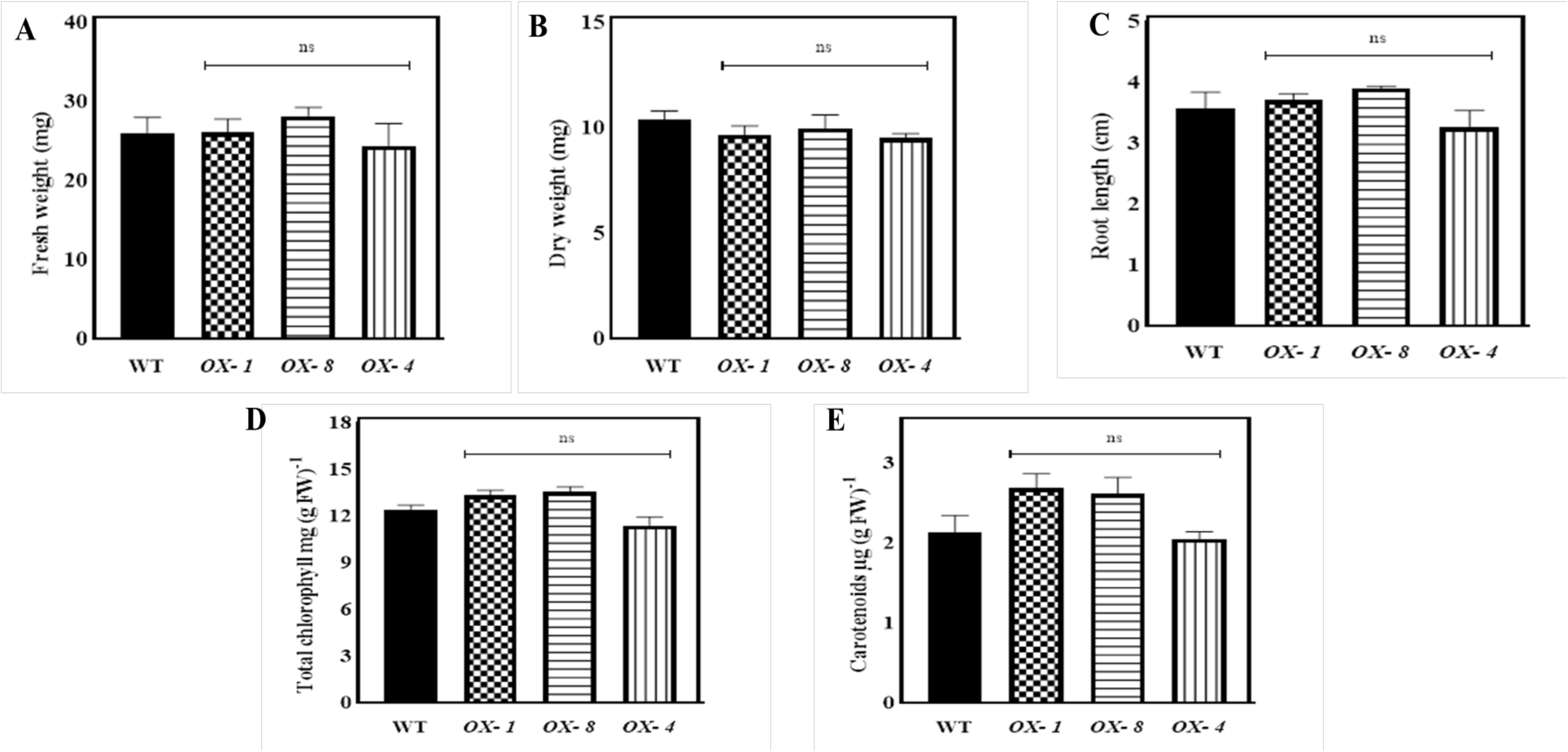
Morphological and physiological parameters of *OsGGCT1* transgenic plants. WT and transgenic lines (*OX- 1,4* and *8*) were grown on MS medium under normal conditions at 22 c under 16h/8h light/dark cycle for 21 days and their morphological (A) fresh weight, (B) dry weight, (C) root length and physiological parameters (D) total Chl and (E) carotenoids were analyzed. The tests were analyzed using one-way ANOVA, where *, **, and *** represent significance and **** high significance at *p*c≤c0.05, 0.01, 0.001, and 0.0001, respectively.

**Supplementary figure 3:**
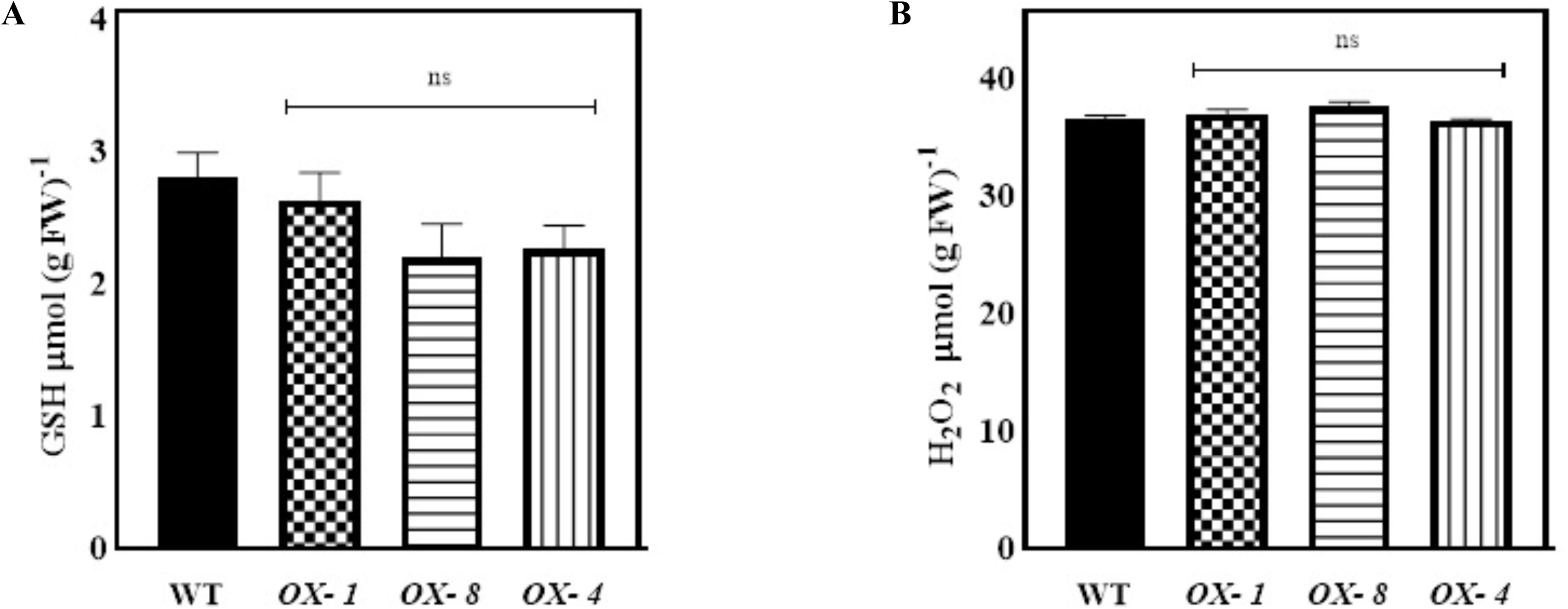
GSH and H_2_O_2_ levels of *OsGGCT1* transgenic plants. WT and transgenic lines (*OX- 1,4* and *8*) were grown on MS medium under normal conditions at 22 c under 16h/8h light/dark cycle for 21 days and their (A) GSH (B) H_2_O_2_ contents were analyzed. The tests were analyzed using one-way ANOVA, where *, **, and *** represent significance and **** high significance at *p*c≤c0.05, 0.01, 0.001, and 0.0001, respectively.

**Supplementary figure 4:**
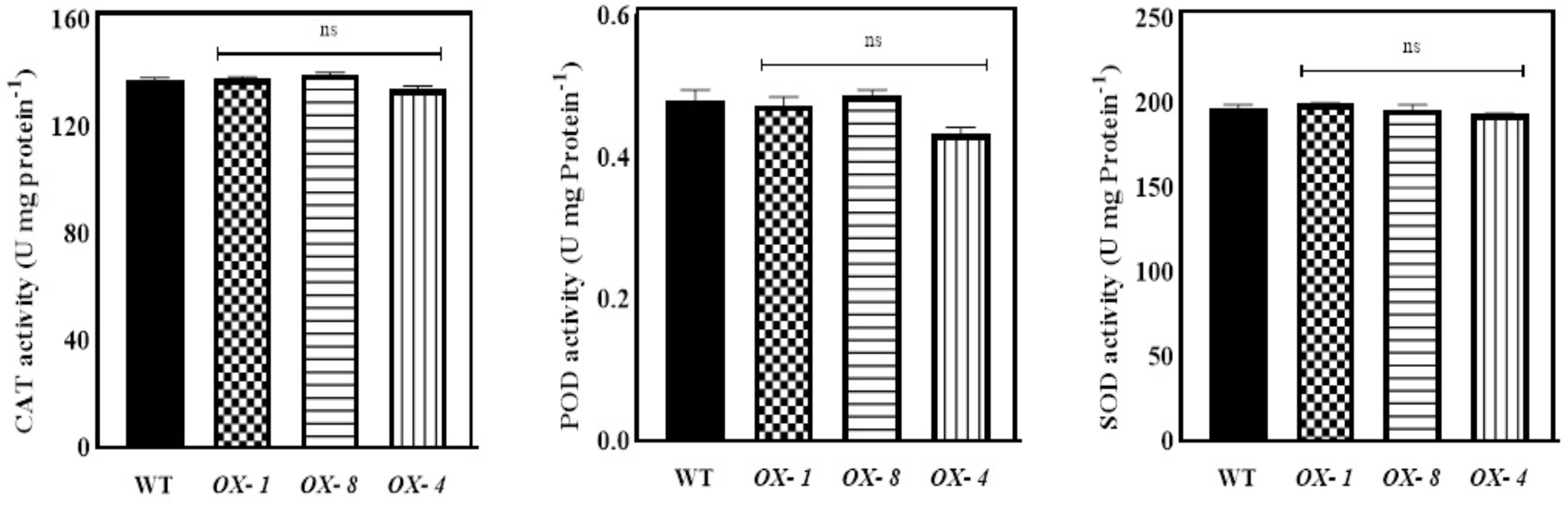
Enzyme activities of *OsGGCT1* transgenic plants. WT and transgenic lines (*OX- 1,4* and *8*) were grown on MS medium under normal conditions at 22 c under 16h/8h light/dark cycle for 21 days and their enzyme activities were analyzed. (A) CAT, (B) POD and (C) SOD. The tests were analyzed using one-way ANOVA, where *, **, and *** represent significance and **** high significance at *p*c≤c0.05, 0.01, 0.001, and 0.0001, respectively

